# Epigenetic regulation of Neuregulin-1 tunes white adipose stem cell differentiation

**DOI:** 10.1101/2020.03.23.004275

**Authors:** Alyssa D. Cordero, Evan C. Callihan, Rana Said, Yasir Alowais, Emily S. Paffhausen, John R. Bracht

**Author notes:** co-first author (Alyssa D. Cordero, Evan C. Callihan). Correspondence: John Bracht.

## Abstract

Expansion of subcutaneous adipose tissue by differentiation of new adipocytes has been linked to improvements in metabolic health. However, an expandability limit has been observed wherein new adipocytes cannot be produced, the existing adipocytes become enlarged (hypertrophic) and lipids spill over into ectopic sites. Inappropriate ectopic storage of these surplus lipids in liver, muscle, and visceral depots has been linked with metabolic dysfunction. Here we show that Neuregulin-1 (NRG1) serves as a regulator of adipogenic differentiation in subcutaneous primary human stem cells. We further demonstrate that DNA methylation modulates NRG1 expression in these cells, and a 3-day exposure of stem cells to a recombinant NRG1 peptide fragment is sufficient to reprogram adipogenic cellular differentiation to higher levels. These results define a novel molecular adipogenic rheostat with potential implications for the expansion of adipose tissue *in vivo*.

## Introduction

Obesity, defined as having a body-mass index (kg/m^2^) of over 30, affects 39.8% of adults in the United States [1], is generally associated with an increased incidence of health problems including type 2 diabetes, cardiovascular disease, and cancer [2]. Obesity is characterized by an increase in percent body fat driven by the expansion of white adipose tissues (WAT). This can occur by an increase in adipocyte size, adipocyte number, and/or accumulation of adipose tissues in differing depots. These different mechanisms of adipose tissue expansion and deposition are associated with differing health outcomes. More specifically, the accumulation of adipose tissue in the visceral regions (intra-body cavity and near vital organs) is associated with an elevated risk of diabetes [3] and cardiovascular disease [4], while subcutaneous lipid storage (within depots just under the skin) is comparatively benign [5,6]. Indeed, the visceral-to-subcutaneous adiposity ratio has been proposed as a predictor of metabolic health [7,8] and a study of obese individuals that transitioned from metabolically healthy to unhealthy obesity showed a corresponding increase in visceral fat [9]. Evidence has begun to accumulate suggesting that adipogenesis-- differentiation of new fat cells--within subcutaneous depots is a key factor in modulating healthy versus unhealthy adipose tissue deposition. The inability of subcutaneous adipose tissue depots to differentiate new fat cells leads to enlargement of existing adipocytes (hypertrophy) and spillover of lipid into ectopic sites including the visceral depots [10–12]. Consistent with this, patients exhibiting visceral adiposity exhibit lower adipogenic potential in subcutaneous depots [13] due in part to senescence of the stem cells [14], suggesting that tissue-specific adipose expandability limits are critical regulators in whole-body metabolic health.

To begin to understand key mechanisms regulating adipose tissue deposition and expandability we utilized Adipose-derived Stem Cells (ASCs). These primary human cells are isolated from subcutaneous adipose tissue lipoaspirates, which are capable of differentiating into multiple lineages, including adipocytes, *in vitro* [15,16]. Because ASCs are derived directly from patient subcutaneous adipose depots, they provide a unique model system to investigate the control of adipocyte differentiation. We demonstrate that epigenetic programming, established *in vivo* prior to isolation from a patient, can be manipulated prior to induction of differentiation *in vitro*, thereby altering NRG1 expression and differentiation behavior. We further show that exposure of stem cells to recombinant NRG1 EGF domain is sufficient to reprogram adipose differentiation apart from epigenetic interference. Since ASCs are obtained as heterogeneous mixtures of cell types [17] we performed the initial experiments in a clonal cell line, but confirmed our findings in raw (heterogeneous) ASCs (raw Processed Lipoaspirates or PLA [15,16]).

Here we show that Neuregulin-1 (NRG1) plays the role of master regulator of WAT differentiation from subcutaneous primary human stem cells, and that it is under epigenetic control. Consistent with its known role in other tissues, our data shows NRG1 regulates formation of new adipocytes from stem cell precursors.

## Methods

### Stem Cell Culture

Human Adipose-Derived Stem Cell (ASC) lines (Table 1) were cultured at 37°C with 5% CO_2_ in Growth Media consisting of Dulbecco’s Modified Eagle’s Medium (DMEM) supplemented with 10% fetal bovine serum (FBS), 1% Penicillin (10,000 u/mL)-Streptomycin (10,000 μg/mL), and 1X Glutamax. Growth Media was changed three times per week.

**Table 1.** Cell lines used in the study.

| Cell Line | BMI | Depot | Sex | Age | Passage at Experiment | Source | Figure |
| --- | --- | --- | --- | --- | --- | --- | --- |
| ASC080414A-derived clonal line = AF2 | 25.1 | Abdomen | F | 39 | p12 | Zen-Bio, Inc. | Fig. 1A, Fig. 2B, Fig. 3, Fig. 4A |
| ASC072709 | 38.0 | Hip | F | 39 | p9 | Zen-Bio, Inc. | Fig. 4B |
| Line 1107 | 27.4 | Abdomen | F | 40 | p8 | DeCicco-Skinner lab (AU) | Fig. 4C |
| ASC012502 | 25.3 | Abdomen | M | 40 | p4 | Zen-Bio, Inc. | Fig. 2C |

### Generation of Clonal ASC Line

Polyclonal ASC culture ASC080414A (Zen-Bio, Inc.) was trypsinized and neutralized in DMEM supplemented with 20% FBS, 1% Penicillin-Streptomycin, and 1X Glutamax, and quantified using a hemocytometer. Cells in suspension were then subjected to serial limiting dilution into a 96-well dish. After 12-24h, wells containing a single cell were identified by inspection and cultured to give rise to monoclonal ASC lines. After 1 week, monoclonal lines were switched back to standard growth media (DMEM with 10% FBS, 1% Penicillin-Streptomycin, 1X Glutamax). Clone F2 demonstrated strong growth, was renamed AF2 (short for ASC080414<u>A-F2</u>) and and was used in epigenetic and differentiation experiments for gene expression analysis.

### Decitabine (DAC) Treatment and Adipose Differentiation

AF2 cells (passage 8) were cultured to approximately 80% confluence in 6-well dishes (3 wells per treatment). 5-aza-2’-deoxycytidine (Decitabine, DAC) was diluted in DMSO to stock concentrations of 5mM and 0.625mM. A total of 0.4μL of each stock solution of DAC was added to respective wells containing cells in 2mL of Growth Media, giving final concentrations of 1μM and 0.125μM. 0.4μL DMSO was added to 3 wells as carrier controls. DAC or DMSO was added once per day for 3 days, with no media change. On the fourth day, one set of replicates (total of 3 replicates per treatment) was harvested for DNA and RNA using ZR-Duet DNA/RNA MiniPrep (Plus) kit (Zymo Research Cat # D7003) following manufacturer’s protocol. Following extraction, RNA samples were further cleaned and concentrated using the RNA Clean & Concentrator kit (Zymo Research Cat # R1013) into 15μL of DNAse/RNase-Free Water as per the manufacturer’s protocol. RNA was sent to the Deep Sequencing Core at Johns Hopkins for Affymetrix Microarray analysis.

Additional DAC or DMSO-treated AF2 replicates (three per treatment) were switched to Adipogenic Media and cultured for 18 days, at which point adipocyte differentiation was quantitated by imaging. The adipogenic differentiation media consisted of DMEM supplemented with 10% FBS, 1% Penicillin Streptomycin, 1X Glutamax, 1.0 μM Dexamethasone, 0.5 mM IBMX (3-isobutyl-1-methylxanthine), 0.2 mM Indomethacin, and 10.0 μM Insulin.

### NRG1 Recombinant Protein Experiment in 6-well dishes

Two recombinant NRG1 isoforms, NRG1-α, Cat #559502 and NRG1-β, Cat #711104 were ordered from BioLegend. NRG1-α (65 amino acids) was diluted in DPBS supplemented with 1% Bovine Serum Albumin (BSA) to 40ng/μL, 8ng/μL, 1.6ng/μL, 0.32ng/μL, and 0.064ng/μL. NRG1-β (65 amino acids) was reconstituted in to 200ng/μL in sterile H_2_O supplemented with 5% Trehalose as recommended by the manufacturer, and serially diluted to the same concentrations as NRG1-α. These stocks provided 1:1000 dilutions, and 2μL of each was added to corresponding wells of a 6-well dish (approximately 80% confluence) along with fresh growth media. For controls, 2μL of 1% BSA or 5% Trehalose were used and 3 wells were left untreated. Stem cells were allowed to grow for 3 days with recombinant NRG1 (or carrier control) to reach 100% confluence and then were switched to adipogenic media (DMEM with 10% FBS, 1% Penicillin Streptomycin, 1X Glutamax, 1.0 μM Dexamethasone, 0.5 mM IBMX (3-isobutyl-1-methylxanthine), 0.2 mM Indomethacin, and 10.0 μM Insulin) and allowed to differentiate for 18 days (without recombinant protein). On day 18, differentiation was quantified by microscopic imaging. For each well, 4 “light” images (showing all the cells present in the field of view) and 4 corresponding “dark” images (using high contrast to highlight lipid droplets characteristic of adipocytes) were taken. The total number of cells, as well as the number of lipid (+) adipocytes, present in an image was determined using ImageJ software.

### Adipocyte differentiation quantitation with Oil Red O in 24-well dishes

NRG1 validation experiments were conducted in 24-well dishes, cultured and treated with the same concentrations of recombinant protein as described for the 6-well dish experiment. As in the 6-well dish experiments, differentiation occurred for 18 days after the switch to adipogenic media.

The differentiation was quantitated with Oil Red O (OrO). Oil Red O (OrO) working solution was prepared: OrO stock solution (generated by mixing 0.3g Oil Red O powder, Santa Cruz Biotech. CAS# 1320-06-5) was dissolved into 100ml of 100% isopropanol. From this stock, a working solution was created by adding 24ml OrO stock to 16ml distilled H_2_O (dH_2_O), which was mixed thoroughly and allowed to stand for 10 minutes prior to filtration through a 0.2μM vacuum filter unit (VWR, Cat # 10040-460) to remove particulates. This working stock is usable for up to 3 hours.

For all experiments, several wells of an unseeded 24-well dish was used as no-cell controls, because of a marked tendency for Or to stick to polystyrene. All differentiated stem cells and the no-cell control wells were washed with DPBS (1ml per well, Gibco Cat# 14190144) and fixed for 30min with 1m of PBS (pH 7.2) supplemented with 4% paraformaldehyde and 1% CaCl_2_. After fixing, cells were washed once with DPBS and stained with 1ml of OrO working solution for 15 minutes. OrO working solution was removed and the wells were washed with dH_2_O three times. To prevent drying out, one dish was aspirated and re-filled at a time; if wells dry out lipid and the OrO signal within the cells is lost. After final wash and aspirate, the OrO signal was eluted by adding 1ml of 100% isopropanol to all wells and gently rocking for 15 minutes (visually confirm elution under the microscope and allow more time if required). After extraction, pipet 100μl of the isopropanol-OrO into a clean 96-well plate; measure each extracted sample in triplicate (three wells in the 96 well plate) as technical replicates. To quantify OrO extraction, measure absorbance at 450 nm with a microplate reader; perform subtraction of average no-cell control signal to correct for background binding to polystyrene.

### NRG1 Isoform Expression Analysis

RNA was extracted and DNAseI-treated from DAC-treated AF2 cells using the ZR-Duet DNA/RNA MiniPrep (Plus) kit (Zymo Research Cat # D7003). This RNA was converted to cDNA using SuperScript III First-Strand Synthesis System (Invitrogen Cat# 18080051) according to manufacturer’s instructions. Quantitative PCR was carried out using PowerSYBR master mix (ThermoFisher Cat# 4367659) utilizing primers specific to the unique first exon of each NRG1 isoform (Type I, Type II, and Type III, STAR methods). Beta-Actin primers obtained from Huang, et al 2011, were used as a normalization control. The qPCR used a 2-step program, with 40 cycle of 95°C denature (30s) and 60°C anneal/extend (60s).

### Bisulfite Conversion and PCR

DNA was isolated from DAC-treated 414A F2 monoclonal cell line, and 250ng of DNA was bisulfite converted per sample using the EZ DNA Methylation-Gold kit (Zymo Research) according to the manufactuer’s protocol. PCR was performed on bisulfite-converted material with “Bisulfite CpG Island III” primers using the following program: (step 1) 94°C 2 min, (step 2) 94°C 30 seconds, (step 3) 49°C 30 seconds, (step 4) 72°C for 30 seconds, repeat steps 2-4 for 40 cycles, (step 5) 72°C for 5 minutes, (step 6) 4°C infinite hold.

## Results

Given that NRG4 is a marker for brown adipose tissue (BAT) [18], we investigated the expression of all four Neuregulin paralogs (NRG1-4) in primary clonal human white adipose-derived stem cells (ASCs) and *in vitro* differentiated adipocytes. While these cells (AF2) express only negligible levels of NRG 2-4, they exhibit detectable levels of NRG1 (Figure 1A). A similar expression pattern was seen in murine cell lines [17] derived from inguinal (subcutaneous, Figure 1B) and perigonadal (visceral, Figure 1C) fat pads [17]. Differentiation into adipocytes did not appreciably alter expression of any neuregulin (Figure 1A). We used an additional marker of WAT, UCP2, which we found in both human and mouse cells as demonstrating these cells are WAT [19], not BAT (BAT preferentially expresses UCP1[20]). Our previously published analysis demonstrates that perilipin-1, leptin, and other genes behave as expected upon adipogenic differentiation in this cell line [21].

**Figure 1.**
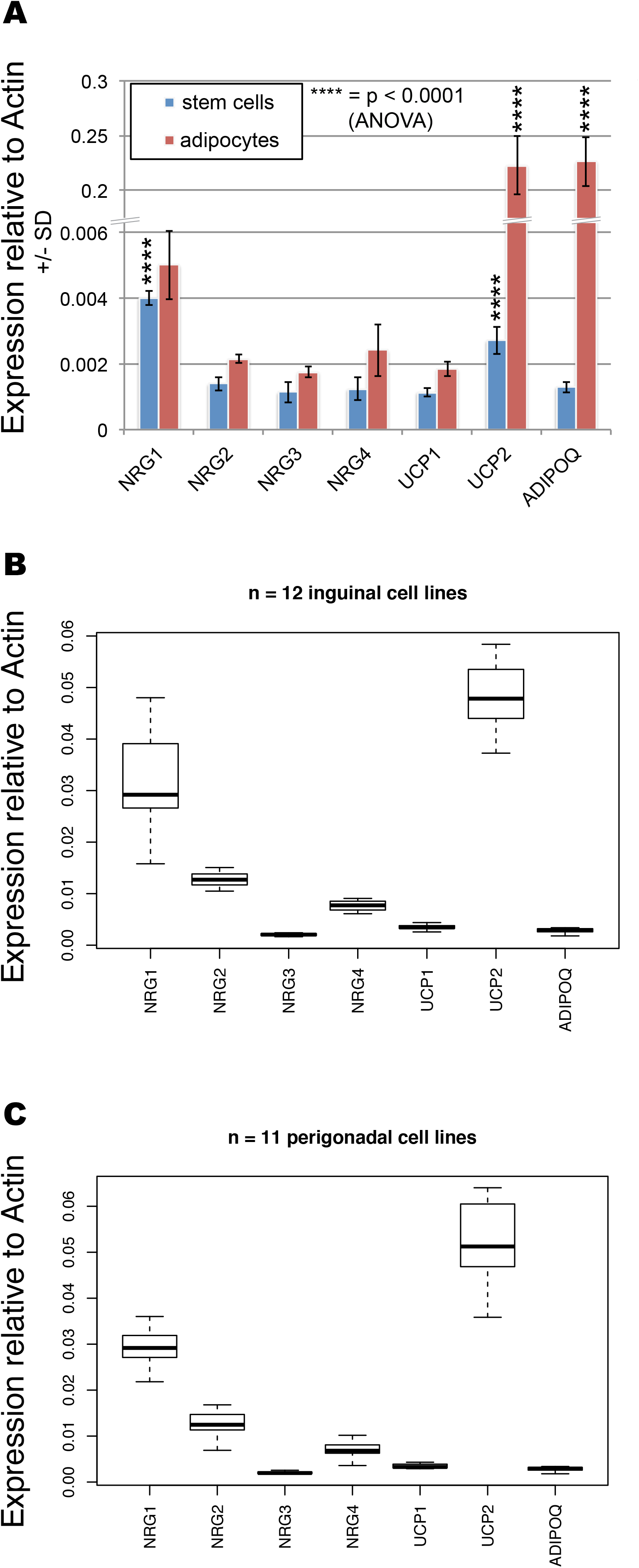
Expression of NRG1-4 and markers of brown (BAT) or white adipose tissue (WAT) in human and murine primary stem cells. A. Human data from clonal AF2 cells (ASCs). NRG1 is expressed preferentially to NRG4 (BAT marker) in both stem cells and adipocytes. UCP1 is a BAT marker and UCP2 is a WAT marker. Adiponectin (ADIPOQ) is shown as a differentiation control. P-values calculated relative to corresponding (differentiated or undifferentiated) NRG4 using 1-way ANOVA and Tukey’s HSD post-hoc test. B. Analysis of gene expression in mouse adipose precursor cell lines isolated from inguinal depots of a male mouse. C. Gene expression in mouse adipose precursor cell lines isolated from perigonadal depots of a male mouse. For both B and C, ANOVA analysis followed by Tukey’s HSD post-hoc test reveals that NRG1 expression is significantly different (p<<0.01) from all other genes. Microarray data from [17]. NRG, Neuregulin; UCP, uncoupling protein; AdipoQ, adiponectin.

Because of NRG1’s well-established role promoting differentiation of stem cells in neuronal [22], retinal [23] and heart [24] tissue, we hypothesized that NRG1 may be critical in defining the differentiation potential of WAT stem cells, possibly through epigenetic regulation. To test for epigenetic control of NRG1 we exposed AF2 cells to the DNA methylation inhibitor 5-aza-2’-deoxycytidine (DAC) and assessed expression of three NRG1 isoforms, each with a unique transcriptional start site leading to expression of a unique 5’ exon (Figure 2A, [25]) by RT-qPCR analysis. While the Type I isoform was mildly induced (~2-fold) by DAC treatment and the Type II isoform was not detected, the Type III isoform exhibited a five-fold and ten-fold up-regulation upon treatment with 0.125 μm and 1 μm DAC respectively (Figure 2B). The same pattern was observed on DAC treatment of raw PLA (Figure 2C) showing that epigenetic regulation of NRG1 Type III isoform may be a general mechanism.

**Figure 2.**
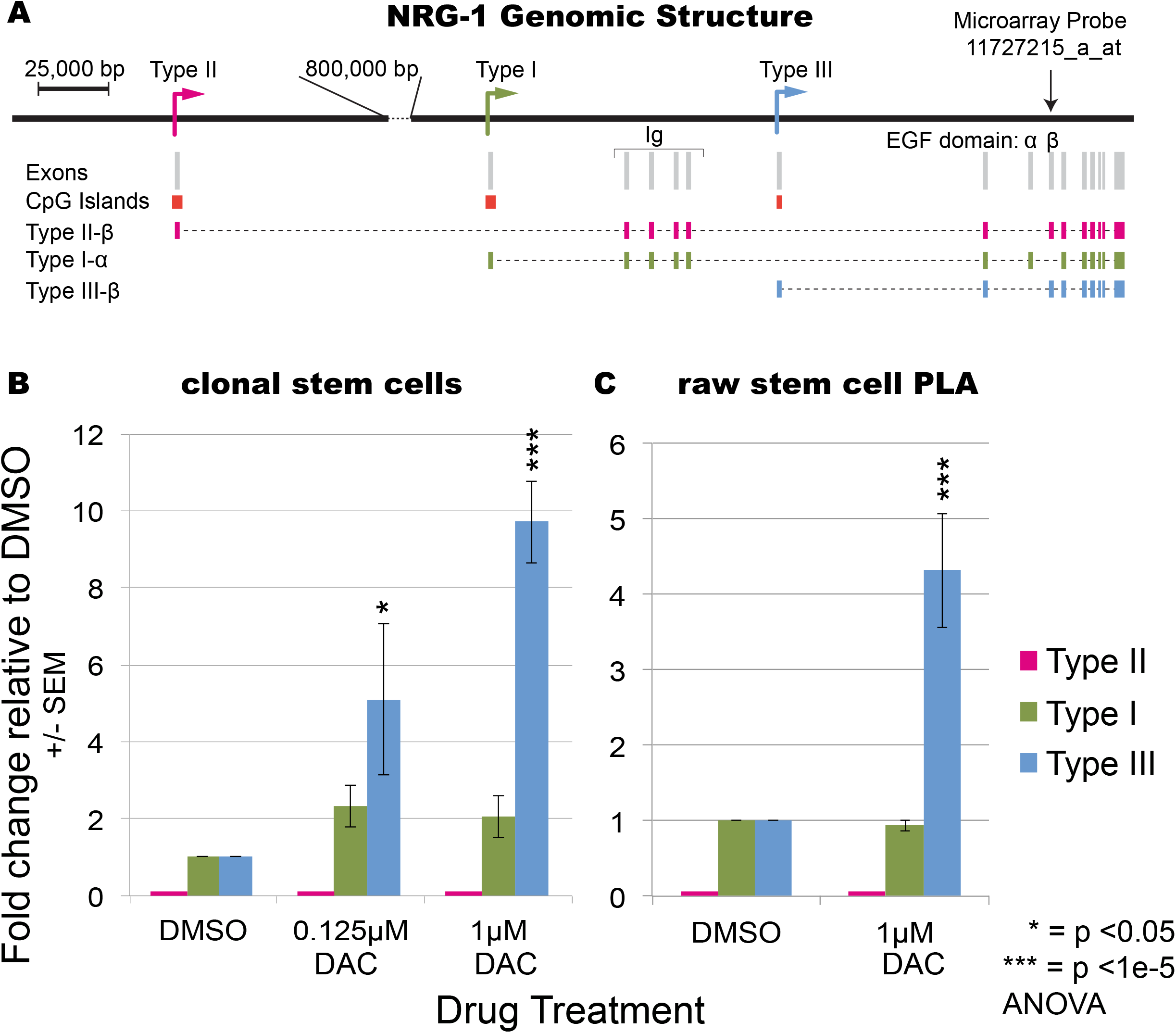
Isoform-specific epigenetic induction of NRG1 expression in primary human ASCs. A. Genomic structure of NRG-1. Types I-III are alternative transcription start sites within the NRG1 locus, each carrying a unique first exon targeted in RT-qPCR. Induction was specifically of Type III isoform was observed (B) in clonal AF2 cells and (C) raw processed lipoaspirate (PLA). Expression of NRG1 is induced with demethylating agent decitabine (DAC). P-values derived from Tukey’s HSD post-hoc test after ANOVA and are relative to corresponding matched DMSO control. Each experiment was performed in triplicate and the fold-change and standard error of the mean are shown.

The increased NRG1 Type III expression induced by DAC treatment correlated with increased cellular differentiation 18 days later in the same experiment (Figure 3A, 3B, 3C). Remarkably, the Pearson’s correlation between adipose differentiation and Type III isoform expression after DAC treatment was 0.989 (Figure 3D). To test whether other genes might be driving this effect, we performed microarray analysis on the DAC-treated AF2 stem-cell RNA. A probe within NRG1 (11727215_a_at, Figure 2A) showed by far the most statistically robust change between DMSO and 1μM DAC (Figure 3E), though it was not the highest fold-change. This probe is in a universal exon so it reports all isoforms together (Figure 2A); by plotting the RT-qPCR data onto the volcano plot this probe may have predominantly reported the Type I isoform’s ~ 2-fold increase (Figure 2B). In contrast, the NRG1 Type III isoform expression increased approximately ten-fold by RT-qPCR (Figure 2B, 3E).

**Figure 3.**
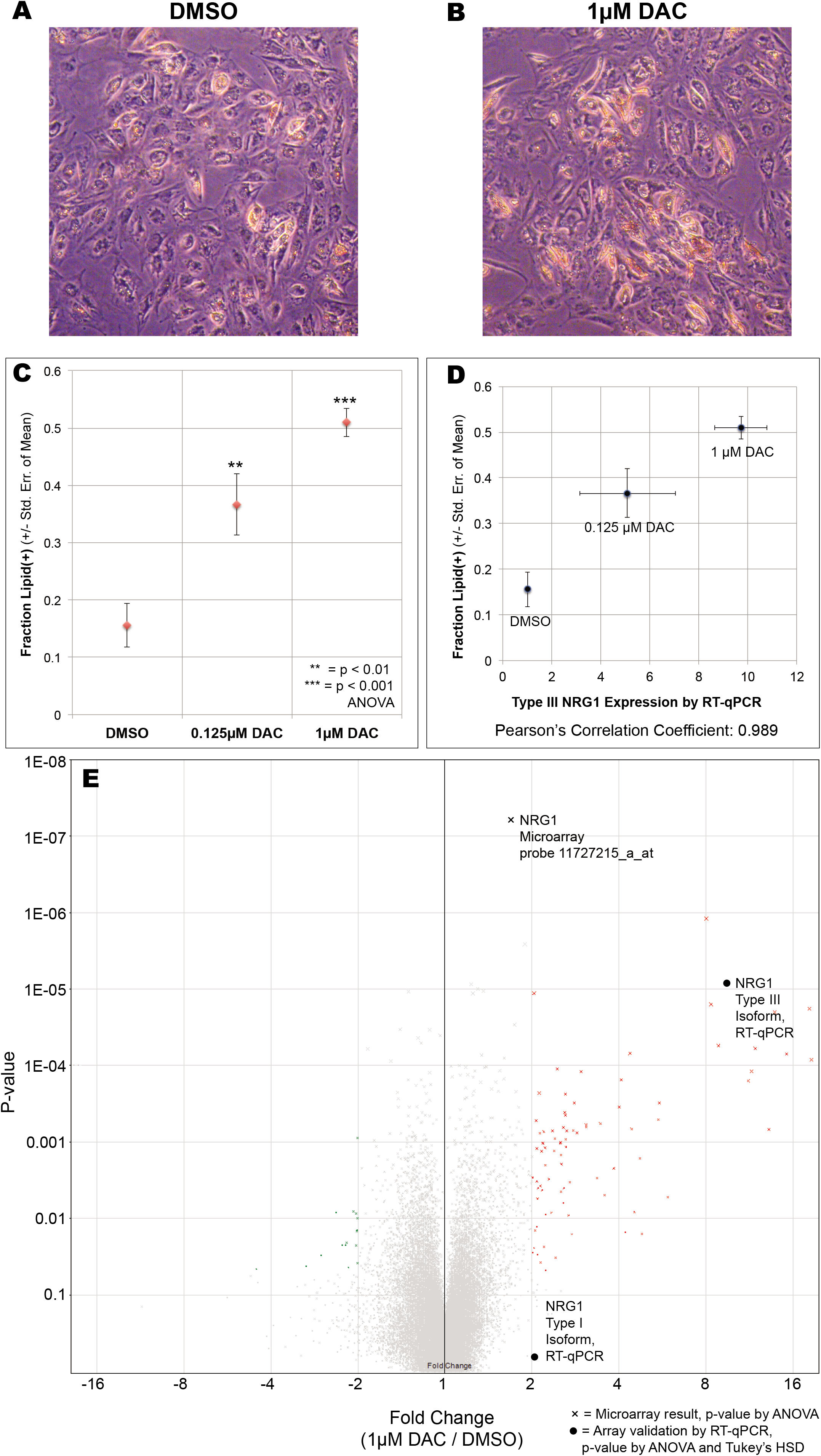
Epigenetic regulation in primary adipose-derived stem cells alters differentiation behavior. **A.** Image of DMSO (control)-treated AF2 cells after differentiation into adipocytes. Accumulated lipid is a straw color on the purple background. **B.** DAC-treated stem cells were differentiated and show increased lipid-positive cells. **C.** Quantitation showing DAC causes increased differentiation. All experiments performed as three biological replicates, with error bars showing S.E.M. **D.** The NRG1 microarray probe (11727215_a_at) correlates with differentiation: Pearson’s correlation coefficient 0.993. **E.** Volcano plot of microarray data, 1.0 μM DAC vs. DMSO (P-values by AOVA). Validation by RT-qPCR is also shown on the volcano plot as filled circles at appropriate fold-change (p-value calculated by ANOVA followed by Tukey’s post-hoc test).

In contrast to NRG1, no other DAC-upregulated genes were compelling candidate regulators of cellular differentiation (Figure 3E, Table 2, Supplemental Table 1). However, the top two significantly upregulated genes by fold-change, Keratin-8 (KRT8) [26] and Metallothionein 1G (MT1G) [27] (Table 2), are known to be epigenetically regulated, specifically through DNA methylation of their promoters, in cancer cell lines. Thus the strongly DAC-upregulated genes from the microarray analysis (Table 2) appear to represent ‘endogenous epigenetic reporters’ whose expression shows how well DAC treatment worked in cell culture.

**Table 2.** All genes at least 4-fold upregulated on 1μM DAC relative to DMSO control. P-values from ANOVA for microarray probe. RT-qPCR validation for NRG1 Type III shown in bold (p-value from Student’s 2-tailed t-test).

| Transcript Cluster ID | Fold Change (linear) (1μM vs. DMSO) | ANOVA p-value (1μM vs. DMSO) | Gene Symbol | Description |
| --- | --- | --- | --- | --- |
| 11752634_x_at | 18.58 | 0.000084 | KRT8 | keratin 8, type II |
| 11758298_x_at | 18.25 | 0.000018 | KRT8 | keratin 8, type II |
| 11756989_x_at | 15.22 | 0.000071 | KRT8 | keratin 8, type II |
| 11758184_x_at | 13.85 | 0.00002 | KRT8 | keratin 8, type II |
| 11717386_s_at | 13.22 | 0.000681 | MT1G | metallothionein 1G |
| 11758188_x_at | 11.85 | 0.00006 | KRT8 | keratin 8, type II |
| 11758301_x_at | 11.51 | 0.000118 | KRT8 | keratin 8, type II |
| 11758183_x_at | 11.23 | 0.000159 | KRT8 | keratin 8, type II |
| **RT-qPCR_NRG1_Type III** | **9.72** | **0.00275 (t-test)** | **NRG1** | **Neuregulin-1** |
| 11727248_a_at | 8.85 | 0.000055 | MYH3 | myosin, heavy chain 3 |
| 11733121_s_at | 8.33 | 0.000016 | DAZL | deleted in azoospermia-like |
| 11715280_s_at | 8.03 | 0.000001 | KRT17 | keratin 17, type I |
| 11756072_s_at | 5.92 | 0.005264 | SAA1 | serum amyloid A1 |
| 11727092_x_at | 5.53 | 0.000309 | IL18 | interleukin 18 |
| 11730408_a_at | 5.48 | 0.00051 | C19orf33 | chromosome 19 open reading frame 33 |
| 11753131_x_at | 4.81 | 0.015946 | TM4SF1 | transmembrane 4 L six family member 1 |
| 11717387_x_at | 4.73 | 0.00164 | MT1G | metallothionein 1G |
| 11753130_at | 4.52 | 0.008248 | TM4SF1 | transmembrane 4 L six family member 1 |
| 11755287_x_at | 4.43 | 0.000674 | KRT8 | keratin 8, type II |
| 11756334_x_at | 4.37 | 0.00007 | ANXA3 | annexin A3 |
| 11753129_a_at | 4.21 | 0.015138 | TM4SF1 | transmembrane 4 L six family member 1 |
| 11724283_a_at | 4.08 | 0.000154 | ANXA3 | annexin A3 |
| 11718347_a_at | 4.01 | 0.000348 | S100P | S100 calcium binding protein P |

The strong DAC-responsiveness of NRG1 type III isoform expression (Figure 2B, 3C) prompted bisulfite-PCR examination of its associated CpG island (Figure 2A) which is only 240 bp long and encodes 16 CpG motifs, making it suitable for bisulfite PCR analysis. Using DNA isolated from the same stem cells experiment as Figure 3, we show that while partial methylation (~ 50% at most CpG sites) could be seen in the DMSO control, it was not decreased upon treatment with 1μm DAC, and if anything the methylation increased (Figure S1). While this is not unprecedented--DAC has been shown to increase methylation in some genomic locations [28]-- it suggests that the epigenetic control point in the genome is unknown. We also cannot rule out the possibility that another epigenetically-regulated transcription factor in turn controls NRG1 type III expression; however from the microarray expression data none of the most up-regulated genes after DAC treatment were transcription factors (Table 2) so this explanation appears less likely.

To test whether NRG1 EGF domain is a driver of increased adipose differentiation, we tested the effect of recombinant NRG1 EGF domains on cultured cells. To mirror the DAC experiment we only exposed stem cells to recombinant protein for 3 days prior to induction of differentiation with adipogenic media (see methods). Subsequent differentiation was performed with non-supplemented adipogenic media--in other words, the cells were only exposed to recombinant protein for their last 3 days as stem cells. After 18 days of adipogenic differentiation we observed a statistically significant near-doubling of adipose differentiated cells in the NRG1-treated culture, recapitulating the effect of DAC (Figure 4A). We also replicated these results in two different ‘raw’ PLA culture of obese (BMI 38, Figure 4B) and overweight (BMI 27.4, Figure 4C) patients to verify the effect is general to more than one cell line or one patient, and does not require clonal lines to be observed.

**Figure 4.**
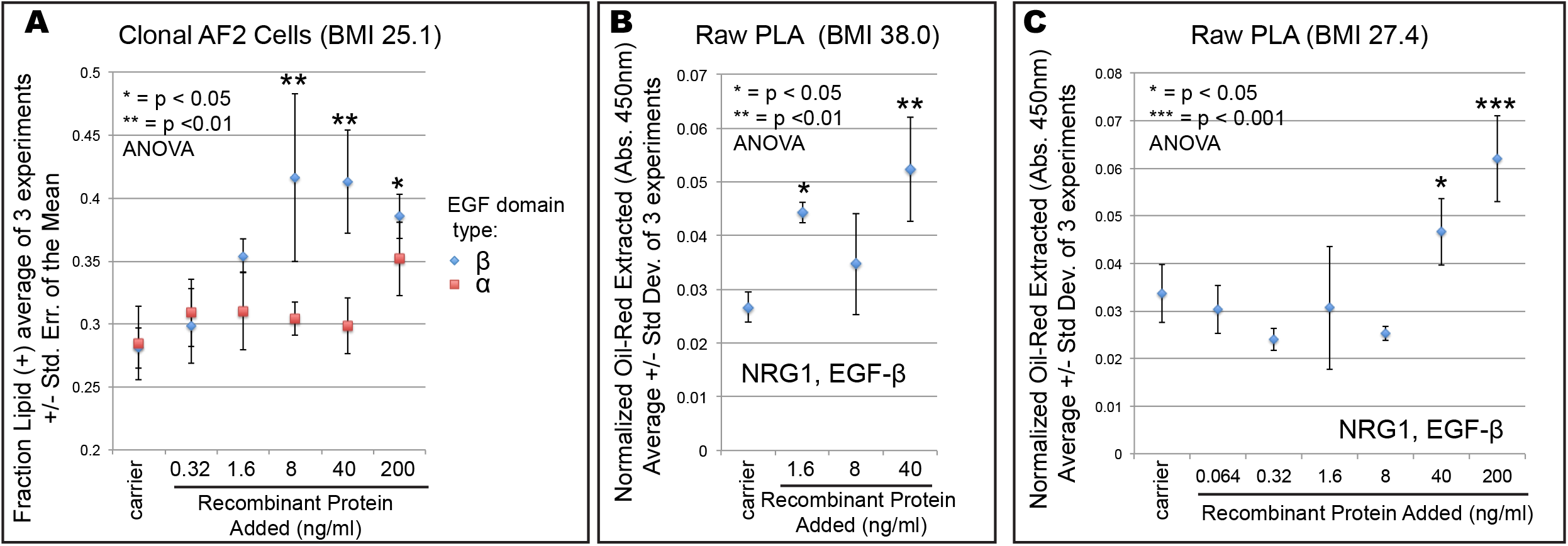
Recombinant NRG1 can re-program stem cells to greater differentiation. **A.** Recombinant NRG1 encompassing either an EGF-α or EGF-β domain was added to the culture media for AF2 clonal cells and adipose differentiation was measured. Shown is the average and standard error of the mean for 3 independent experiments, each with 3 replicates. **B.** Validation with recombinant NRG1 EGF-β in raw PLA (donor BMI = 38.0). Plotted is the average, with error bars representing +/− one standard deviation of 3 replicates. Asterisks represent p-values of 1-way ANOVA relative to the carrier control. **C.** Validation of recombinant NRG1 EGF-β in raw PLA (donor BMI = 27.4). Plotted is the average, with error bars representing +/− one standard deviation of 3 replicates. Asterisks represent p-values of 1-way ANOVA relative to the carrier control.

## Discussion

Neuregulin-4 (NRG4) is a brown adipose marker, identified by transcriptomic analysis of ‘browned’ fat, and may function as an adipokine signal from BAT to neurons [18]. While we find NRG4 is expressed at negligible levels in *in vitro* primary human WAT stem cell culture, we find NRG1 is more highly expressed, both in human (Figure 1A) and mouse (Figure 1B, 1C). Furthermore, NRG1 is epigenetically inducible in human WAT stem cells (Figure 2B, 2C), suggesting it is the WAT-specific neuregulin, a natural counterpart to BAT-specific NRG4. Our analysis of epigenetically inducible NRG1 isoforms showed that a membrane-bound isoform (Type III) is most strongly inducible, suggesting that its role may be more autocrine / juxtacrine than endocrine, potentially acting locally to modulate stem cell differentiation levels within the tissue itself. Because ASCs are primary human cells and tested within 4-12 passages of isolation (Table 1), they retain the patient-encoded epigenetic information and provide a remarkable model system to study adipose-relevant biology. Through a series of experiments we demonstrate epigenetic regulation of NRG1 by DNA methylation regulates the differentiation of WAT stem cells *in vitro*.

This role is consistent with the documented function of NRG1 in other tissues. As already mentioned briefly, in neurons, NRG1 promotes neuronal cell differentiation in the cerebral cortex [22] and retina [23], both *in vivo*, and promotes neuronal differentiation *in vitro* [29]. Similarly, in the heart, NRG1 promotes differentiation of cardiomyocytes from their stem cell progenitors both *in vivo* [24] and *in vitro* [30] and for this reason NRG1 has been successfully tested in clinical trials for heart failure [31,32]. Our results are the first to identify the same differentiation-promoting function of NRG1 within primary adipose-derived stem cell culture.

We hypothesize that the epigenetic control of NRG1 may constitute an intrinsic mechanism--a molecular rheostat--limiting the expansion of subcutaneous WAT depots. In turn, the limits of adipose expansion lead to negative metabolic consequences via ectopic lipid deposition in muscle, liver, or visceral adipose depots [10,11]. Here we suggest that this limiting mechanism is in fact a stem-cell intrinsic epigenetic mechanism of regulation acting through NRG1 as a master regulator.

The identification of NRG1 as a key regulator of adipose expansion may provide a novel therapy for obesity. Wild-type mice treated with injections of recombinant NRG1 display metabolic benefits including lowered bodyweight and reduction in percent body fat relative to controls [33]. NRG1 also improved cardiovascular function and attenuated nephropathy in an *apoE* mutant mouse model [34]. In an obese mouse model (the *db/db* leptin receptor mutant), NRG1 injections improved glucose tolerance [35]. Therefore NRG1 is a candidate for treating metabolic syndrome. When NRG1 was used in clinical trials for human heart failure [31,32] no effect on bodyweight was noted; however, the human therapies were brief, 10 - 11 days, whereas the mouse studies took 8 weeks. Despite multiple studies of systemic administration of recombinant NRG1 in rodents, all of which demonstrated positive metabolic effects [33–37], no study has examined its role in adipose expansion directly. This is even more surprising given that NRG1 administration in mice caused a dramatic spike in leptin, an adipocyte-secreted hormone [33]. Elevated levels of leptin in mouse models promotes reductions in body weight through its effects on the leptin receptor in critical brain regions that regulate food intake and energy expenditure [38].

The dosages of recombinant NRG1 that were active in our experiments are physiologically relevant. Human blood contains variable levels of circulating NRG1 EGF-β isoform, regulated by physical fitness, from 2.6 to 473 ng/ml [39,40], and within the range of our observed *in vitro* activity (Figure 4A,B,C). Even correcting for the size differences between the tested 65aa recombinant fragment and full-length NRG1 (40kDa, [40]) the active concentrations we uncovered are within the physiological range. We also note that the Type III isoform is predicted to remain anchored to the membrane [25] where its effective concentration may be very high for cells in contact with each other in a juxtacrine / autocrine mechanism.

Our data reveal 3-day incubation of recombinant NRG1 β-EGF with ASC is sufficient to reprogram differentiation levels up to 18 days later, without continuous NRG1 exposure. While there is some endogenous bovine NRG1 in the serum used in the culture media, this was accounted for by measuring differentiation of carrier-only controls which were exposed to the same batch of cell culture media.

Our findings also have implications for experimental design. In this study we show that it is possible to treat cells with a non-specific epigenetic agent (decitabine) and to uncover a specific gene (NRG1) by incorporating phenotypic behavior (adipocyte differentiation) as an additional readout. Not only was the NRG1 probe upregulated, but its expression (in stem cells) always tracked very closely with differentiation behavior over two weeks later (Pearson’s correlation 0.989, Figure 3D). This also highlights the value of using clonal cells, minimizing epigenetic noise and cellular heterogeneity of raw PLA [17], and quantitatively evaluating gene expression relative to behavior.

An important future question is to identify the receptor of the NRG1 ligand. Strong candidates are the ErbB family of proteins known to function as NRG1 receptors in other tissues including the brain [41,42]. Interestingly, ErbB gene 4 (ERBB4) has been identified in GWAS studies of both human obesity [43] and diabetic kidney disease [44,45]. These data add a genetic component to the epigenetics of NRG1 and highlight the importance of the NRG1-ErbB signaling axis in human obesity and metabolic health.

The genomic locus where DNA methylation regulates NRG1 expression remains undefined. We found that DNA methylation of the most obvious epigenetic regulatory site—the CpG island just upstream of the Type III promoter (Figure 2A)—is not decreased in the DAC-treated cells (Figure S1). Given the complexity of 3D genome architecture [46], DNA methylation can regulate expression of genes located at large genomic distances. Another possibility is that another gene is upregulated by the epigenetic mechanism directly, and acts as a transcription factor for the NRG1 Type III isoform, though our microarray analysis suggests this is not the case (Table 2). Either way, the epigenetic regulation appears to act at a distance (whether mediated by another gene or not) and the precise epigenetic control point remains to be discovered, a focus of future studies.

Our work is an important contribution to the epigenetic regulation of adipogenesis *in vitro* because we worked in primary human subcutaneous adipose-derived cells. Much previous work utilized 3T3-L1 cells, which are of an unknown lineage from mouse, not human, and are a transformed cell line, not primary cultures [47]. Nonetheless, epigenetic regulation of adipogenic differentiation has been documented in several 3T3-L1 studies [48,49].

Within the human ASC literature, one study demonstrated that inhibiting DNA methylation activates mir193b, thereby promoting adipogenesis of human ASCs [50] suggesting a non-coding RNA signaling axis that may synergize with NRG1-ErbB to promote differentiation. Another study showed that azacitidine treatment inhibits differentiation of ASCs [51], but in that study stem cells were isolated by bariatric and dermolipectomy surgery, not from subcutaneous lipoaspirate, raising the possibility that they represent a different depot (possibly visceral if obtained from bariatric patients). Visceral and subcutaneous depots behave differently and the interplay between visceral and subcutaneous depots may be metabolically critical [13,52]. Another study used bone-marrow derived mesenchymal stem cells (MSCs) to show decitabine (DAC) inhibits adipose differentiation, but these are different cells, from a different depot, than lipoaspirate-derived ASCs [53]. All these studies highlight the diversity of cellular responses to epigenetic modifying compounds, which is particularly important to keep in mind given that ASCs actually represent a heterogeneous mixture [17] and may contain individual cell types that respond differently to epigenetic modulator compounds. We hypothesize that some individual cell lineages within PLA may be ‘positive responders’ and some may be ‘negative responders’ but if they cross regulate each other the overall effect on differentiation may be quite complex and unpredictable or variable from PLA to PLA. This is particularly true as ratios of positive responders and negative responders may even be variant across time within the same culture. We therefore performed most of our work with a clonal line, AF2, which yielded clear results; we then confirmed the findings in raw PLA cultures. It is important to distinguish patient-to-patient variability from intra-patient heterogeneity; even passage numbers affect cellular behavior [54].

While circulating NRG1 has previously been shown to act directly on the central nervous system [22,55], liver [36,55], skeletal muscle [56], and cardiac muscle [24] here we propose an additional site of action within adipose tissues. We present a model that Type III NRG1 is epigenetically regulated and endogenously produced within adipose as a molecular rheostat controlling adipose expandability. In this model, the function of NRG1 is identical to previously published roles positively regulating stem-cell differentiation [22,24]. While this model was developed from *in vitro* data, it explains the dramatic rise in leptin (a primarily adipocyte-secreted hormone) upon administration of NRG1 to wild-type mice [33]; it also may help account for the positive metabolic effects of NRG1 administration to rats [36], hamsters [37] and an obese mouse model [35]. An important direction for future research is to test the model in a rodent by administering NRG1 protein and testing directly for increased adipose expandability, with the ultimate goal of developing therapies for human metabolic syndrome.

## Acknowledgements

The authors wish to thank Dr. Deborah Clegg for valuable comments on the manuscript. The authors also acknowledge Dr. Kathleen DeCicco-Skinner for sharing ASC line 1107 and for providing invaluable help in cell culture for ASCs. This work was supported by two AU Faculty Research Support Grants, one to J.R.B. and another to both J.R.B. and Dr. DeCicco-Skinner. This work was also supported by NIH grant 1K22CA184297 to J.R.B.

## Author Contributions

E.C.C. performed DAC treatment for microarray analysis, NRG1 isoform expression analysis by RT-qPCR, and NRG1 testing with recombinant peptide. R.S. and Y.A. performed stem cell culture experiments supporting the project. E.S.P. performed DAC treatment of stem cells. A.D.C. wrote the manuscript with J.R.B who supervised all work on the project and designed the experiments.

---

**Figure S1.**
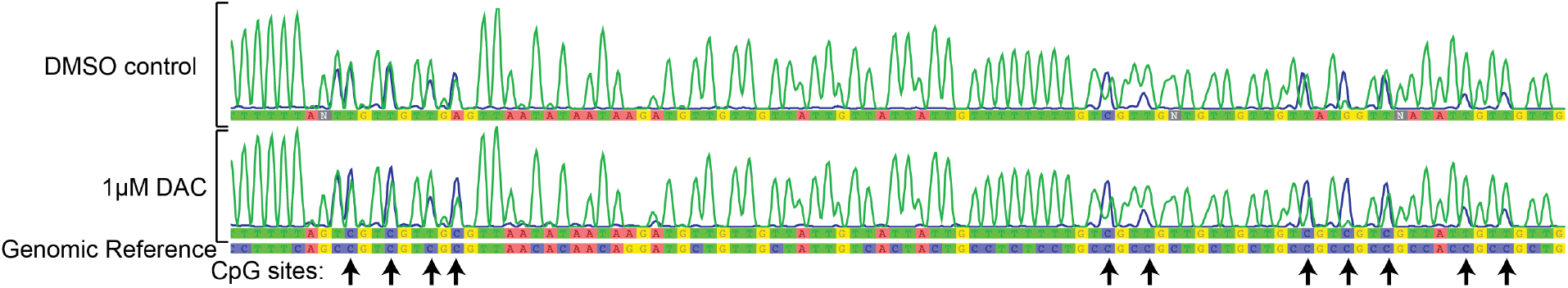
Bisulfate PCR analysis of NRG1 Type III CpG island. Bisulfite treatment causes cytosine (C, blue) to sequence as thymidine (T, green). Methylation is detected as retention of cytosine (blue traces) relative to T (green traces) on chromatograms (for clarity, G and A traces omitted). Note that 1 μm DAC does not demethylate this locus. Arrows mark CpG sites where methylation can occur.

**Supplemental Table 1.**
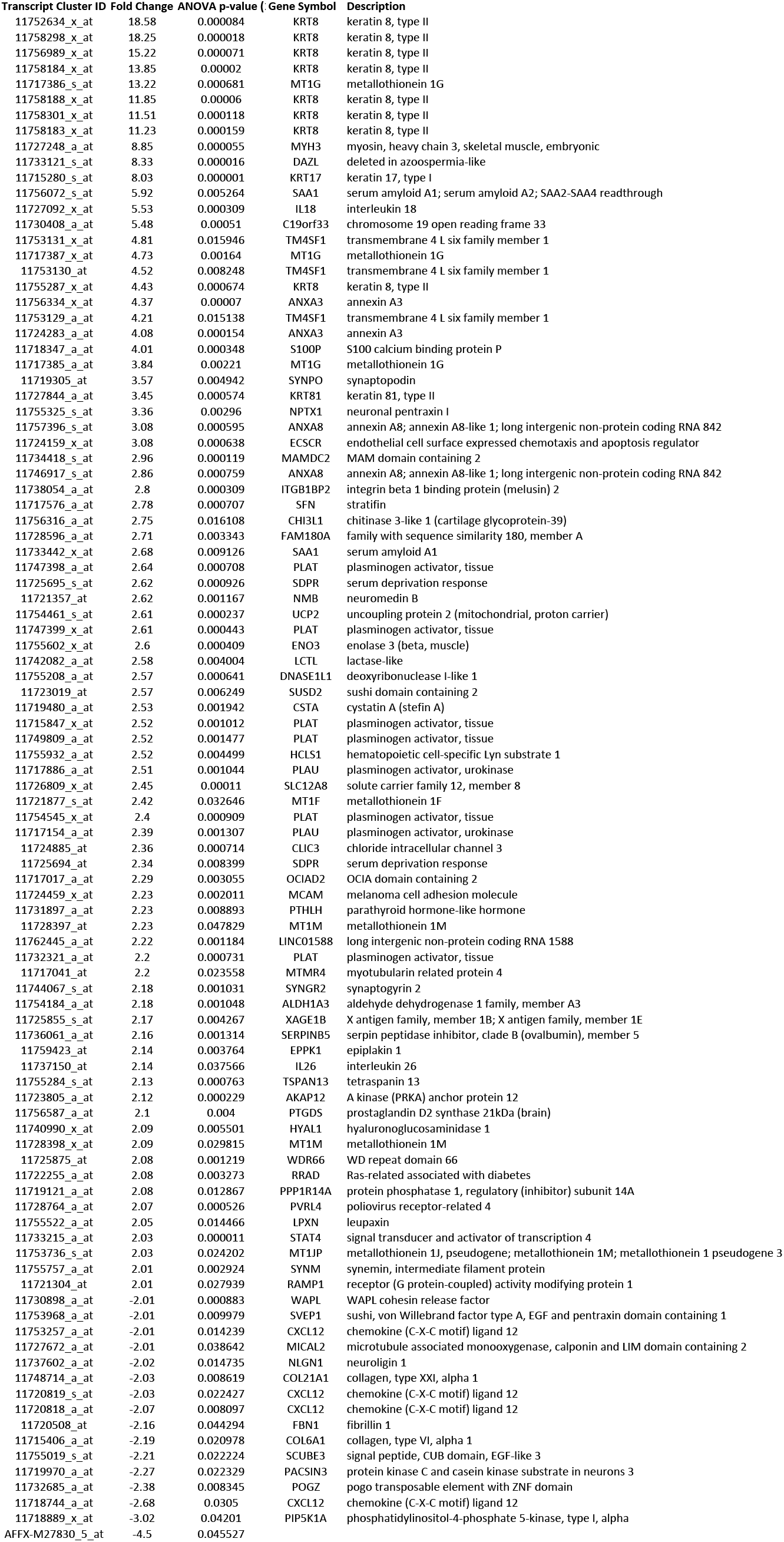
All genes significantly upregulated or downregulated by DAC in volcano plot.

